# CRISPR-directed mitotic recombination enables genetic mapping without crosses

**DOI:** 10.1101/040428

**Authors:** Meru J. Sadhu, Joshua S. Bloom, Laura Day, Leonid Kruglyak

**Affiliations:** Department of Human Genetics, Department of Biological Chemistry, and Howard Hughes Medical Institute, University of California, Los Angeles, Los Angeles, CA 90095, USA

## Abstract

Linkage and association studies have mapped thousands of genomic regions that contribute to phenotypic variation, but narrowing these regions to the underlying causal genes and variants has proven much more challenging. Resolution of genetic mapping is limited by the recombination rate. We developed a method that uses CRISPR to build mapping panels with targeted recombination events. We tested the method by generating a panel with recombination events spaced along a yeast chromosome arm, mapping trait variation, and then targeting a high density of recombination events to the region of interest. Using this approach, we fine-mapped manganese sensitivity to a single polymorphism in the transporter Pmr1. Targeting recombination events to regions of interest allows us to rapidly and systematically identify causal variants underlying trait differences.

---

Identification of DNA sequence differences that underlie trait variation is a central goal of modern genetic research. The primary tools for connecting genotype and phenotype are linkage and association studies. In these studies, co-inheritance of genetic markers with the trait of interest in large panels of individuals is used to localize variants that influence the trait to specific regions of the genome. The localization relies on meiotic recombination events that break up linkage between markers on a chromosome. Therefore, the spatial resolution of genetic mapping is limited by the recombination rate. In practice, the recombination rate in most settings is too low to resolve mapped regions to individual genes, much less to specific variants within genes. Increasing mapping resolution requires construction of ever-larger panels of individuals and/or additional generations of recombination, and these approaches are laborious to the point of often being impractical. As a consequence, the genes and variants underlying trait variation remain unidentified for the vast majority of regions implicated by linkage or association mapping.

To address this problem, we have devised a new method for genetic mapping that precisely targets recombination events to regions of interest. The method uses recombination events that occur during mitosis rather than meiosis. Rare mitotic recombination events occur naturally when a chromosomal double strand break (DSB) is repaired by homologous recombination (HR) that leads to the formation of a recombined chromosome (Yin and Petes 2013). In a heterozygous individual, cell division can then generate daughter cells with a new genotype that is completely homozygous from the recombination site to the telomere and unchanged heterozygous everywhere else (Fig. 1A); such an event is termed “loss of heterozygosity” (LOH). Individuals with LOH events at various locations in the genome have been used to construct a genetic map (Henson *et al*. 1991), and this and related approaches (Laureau *et al*. 2016) can, in principle, be used to map the genetic basis of trait variation (Fig. 1B). However, this approach has been limited in practice by the very low frequency of natural mitotic recombination events.

**Fig. 1:**
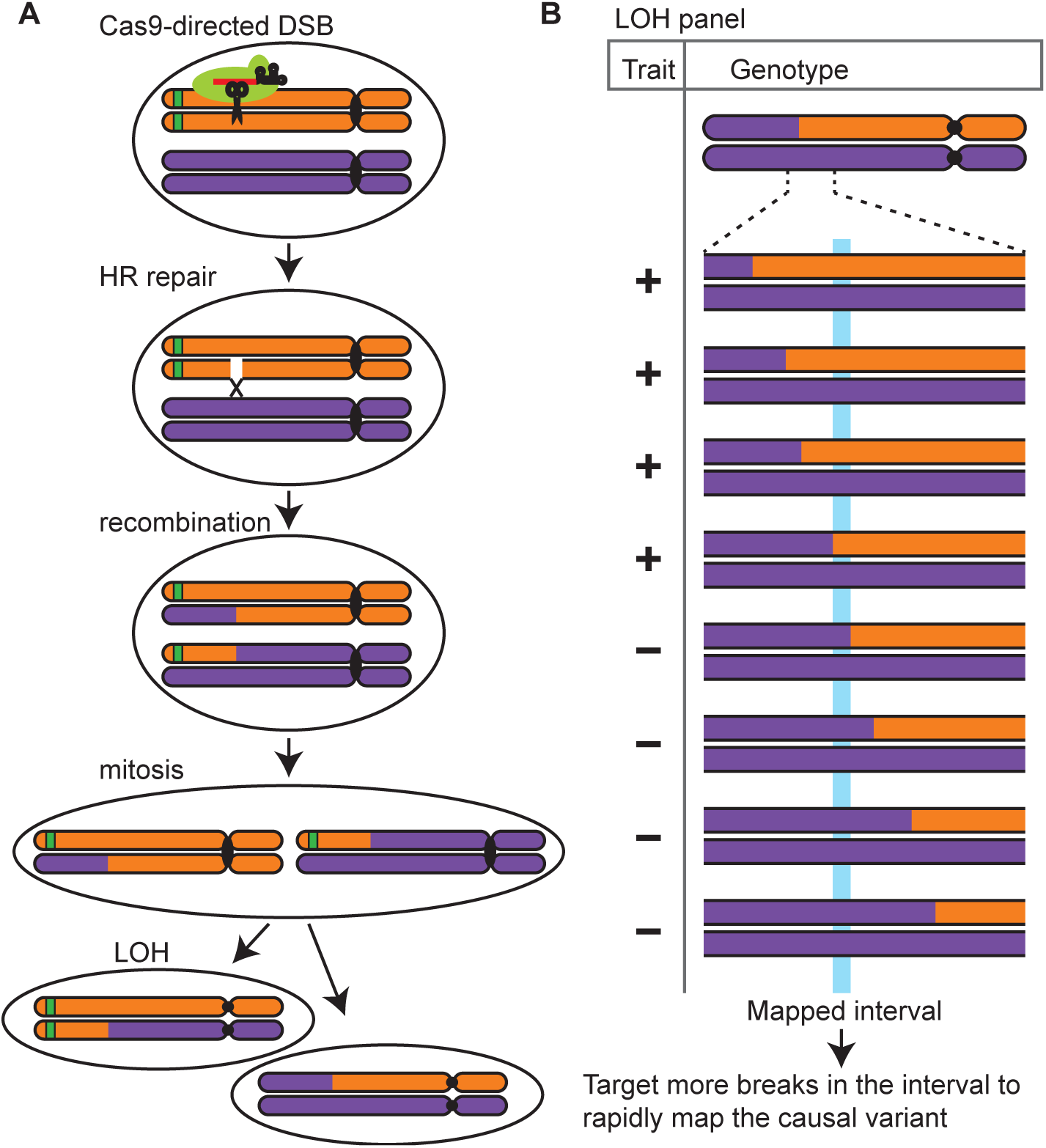
DSBs generated by Cas9 in diploid mitotic cells can lead to mitotic recombination and loss of heterozygosity (LOH). (A) LOH can result from repair following a doublestrand break (DSB) in mitotically dividing cells, which is generated by CRISPR (the *Streptococcus pyogenes* Cas9 protein is depicted as a green cartoon). Individuals with LOH events are isolated via the loss of a heterozygous dominant marker, denoted with a green bar. (B) By measuring trait values in a panel of individuals with LOH events distributed across a region of interest, we can map genetic variants that contribute to trait variation. The process can be iterated to increase mapping resolution.

We have leveraged the CRISPR-Cas9 system to produce targeted mitotic recombination events at high frequency and at any desired location, allowing facile construction of LOH-based mapping panels. In the CRISPR (clustered, regularly interspaced, short palindromic repeats) system, the endonuclease Cas9 creates a DSB at a site specified by the targeting sequence of a bound guide RNA (gRNA) (Doudna and Charpentier 2014). Successful cutting requires the targeted sequence to be followed by an invariant protospacer-adjacent motif (PAM). In a heterozygous diploid individual, an LOH event can be generated by cutting only one chromosome, leaving its homolog intact to serve as a template for repair by HR. This is accomplished by using polymorphic heterozygous PAM sites.

To demonstrate that LOH events can be targeted to precise loci using CRISPR, we designed 95 gRNAs targeting the bacterial *Streptococcus pyogenes* Cas9 to sites distributed across the left arm of the yeast *Saccharomyces cerevisiae* chromosome 7 (Chr 7L). The gRNAs targeted heterozygous sites in a diploid yeast strain generated by crossing a lab strain (BY) and a vineyard strain (RM), using PAMs polymorphic between the two strains. After cutting, repair, and mitosis, cells in which the DSB repair led to an LOH event were isolated by fluorescence-activated cell sorting (FACS) through their loss of a telomere-proximal green fluorescent protein (GFP) gene. We picked approximately four GFP(-) lines per targeted site, for a total of 384 lines. Whole-genome sequencing demonstrated that CRISPR-induced recombination was highly effective, with LOH events in more than 95% of lines and few off-target effects (Supplementary Methods). 75% of LOH recombination events occurred within 20 kb of the targeted site (Fig. 2A), consistent with previous measurements of LOH gene conversion tract length (St. Charles and Petes 2013). LOH events were generated at sites across the entire targeted chromosome arm (Fig. 2A), demonstrating that our method is not limited to certain genomic contexts.

**Fig. 2:**
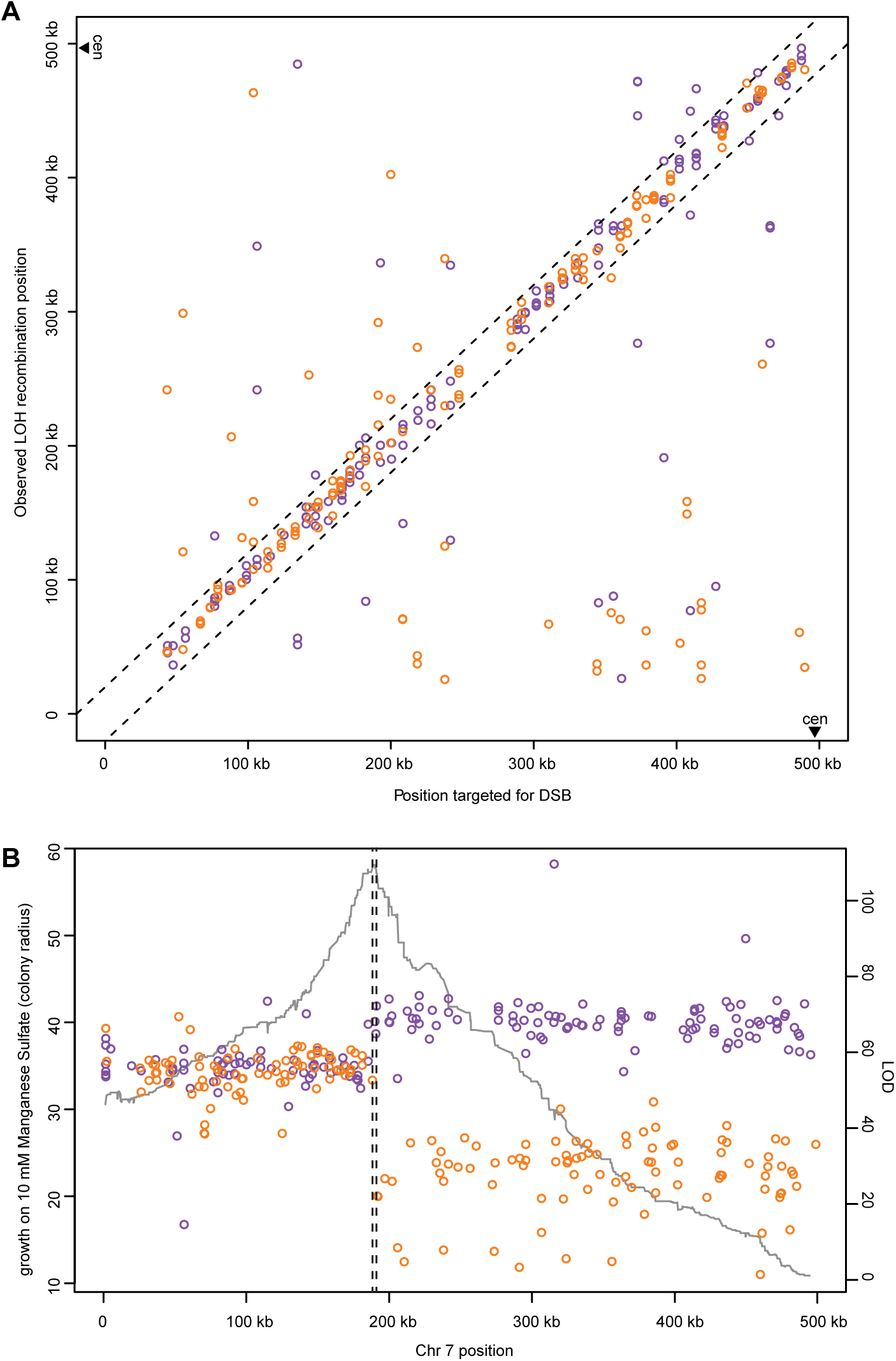
LOH events generated at sites across a chromosome arm mapped manganese sensitivity. (A) For each individual in the panel with a Chr 7L recombination event, the site of its recombination event is plotted against the site targeted for DSB formation in that individual. Individuals targeted to gain BY and RM homozygosity are plotted in orange and purple, respectively. The dashed lines enclose individuals with recombination events within 20 kb of the targeted site. (B) Sensitivity to manganese vs. observed LOH recombination location. For each individual in the Chr 7L panel, the site of the LOH recombination event is plotted against manganese sensitivity, measured as colony radius after growth on 10 mM manganese sulfate plates. Orange and purple points denote individuals that are homozygous BY and RM to the left of their recombination events, respectively. (All individuals are heterozygous BY/RM to the right of their recombination events.) The gray line plots the LOD score by position along Chr 7L for manganese sensitivity. Dashed vertical lines denote the QTL support interval.

We next used the LOH panel to map quantitative traits to loci on Chr 7L. We measured growth of the 384 LOH lines in 12 different conditions, chosen because we previously mapped quantitative trait loci (QTLs) for growth in these conditions to Chr 7L (Bloom *et al*. 2013). In parallel, we measured growth of 768 segregants from a cross between BY and RM. One of the traits, growth on 10 mM manganese sulfate, mapped to a large-effect QTL with a maximum logarithm-of-odds (LOD) score of 109.4 in the LOH panel (Fig. 2B). The confidence interval obtained with the 384 LOH lines overlapped with and was narrower (2.9 kb) than that obtained with 768 segregants (3.9 kb). The LOH-based interval contained two genes and 12 polymorphisms between BY and RM. We identified concordant QTLs of smaller effect in the two panels for eight other traits (fig. S1). Two traits mapped to a QTL of small effect in just one panel, likely due to low statistical power (fig. S2). One trait lacked a Chr 7L QTL in both panels.

To rapidly fine-map the causal variant for manganese sensitivity, we generated a second panel of LOH lines whose recombination events were all targeted to the mapped manganese sensitivity interval. We took advantage of the fact that LOH gene conversion tracts vary in length, which means that in different individuals, DSBs generated by the same gRNA can lead to slightly different LOH crossover sites, typically within 10 kb of the DSB (St. Charles and Petes 2013). We isolated 358 GFP(-) lines generated with three gRNAs targeting sites near the mapped interval. Sequencing revealed that 46 lines (13.1%) had a recombination event within the 2.9 kb QTL interval; together, the recombination events separated almost all the variants in the interval (Fig. 3A). In contrast, only 0.7% of segregants had recombination events in the interval (Bloom *et al*. 2013). To obtain a comparable number of recombination events at this locus by random meiotic segregation, a segregant panel would require more than 7,500 lines. Thus, with targeted LOH events, we can generate very strong mitotic recombination hotspots at any region of interest (fig. S3).

**Fig. 3:**
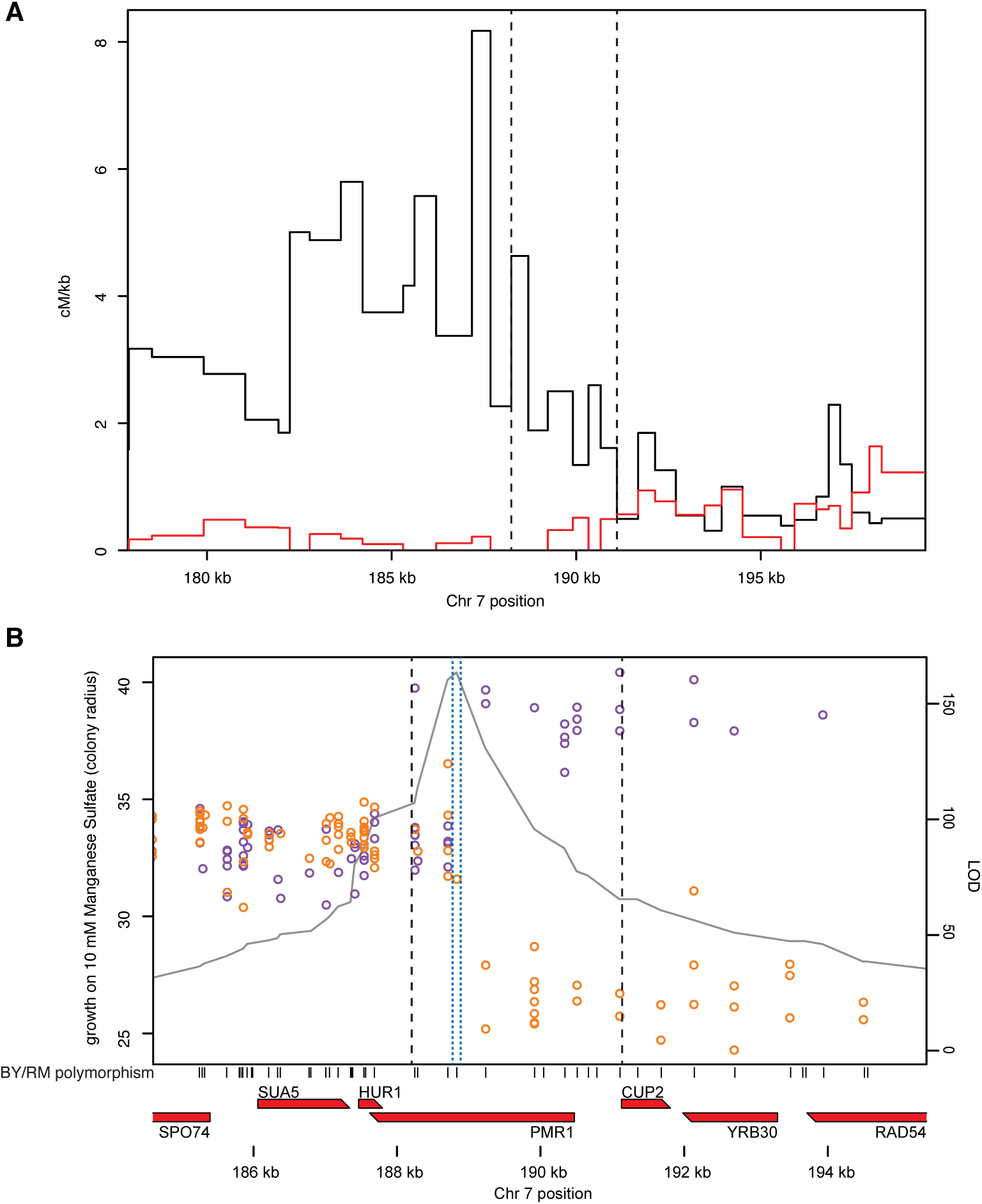
Targeted high-resolution mapping of manganese sensitivity. (A) Ratio of recombination rate (in centimorgans; cM) to physical distance (in kilobases; kb) near the manganese sensitivity QTL, for the manganese fine-mapping LOH panel (black line) and a segregant panel (red line) (Bloom *et al*. 2013). The ratio is plotted for every interval between adjacent BY/RM polymorphisms that are at least 300 bp apart. The fine-mapping panel contains recombination events between all such pairs of polymorphisms in the interval, as the ratio does not drop to zero. The 2.9 kb QTL interval is denoted with dashed lines. (B) Recombination sites of individuals in the fine-mapping panel plotted against their manganese sensitivity, as in Figure 2B, near the manganese sensitivity QTL. Dashed blue lines denote the QTL support interval for the fine-mapping panel and dashed black lines denote the QTL support interval for the whole-Chr 7L panel. Shown below the plot are all BY/RM polymorphisms in the region (black bars), as well as all open reading frames (red lines).

**Fig. 4:**
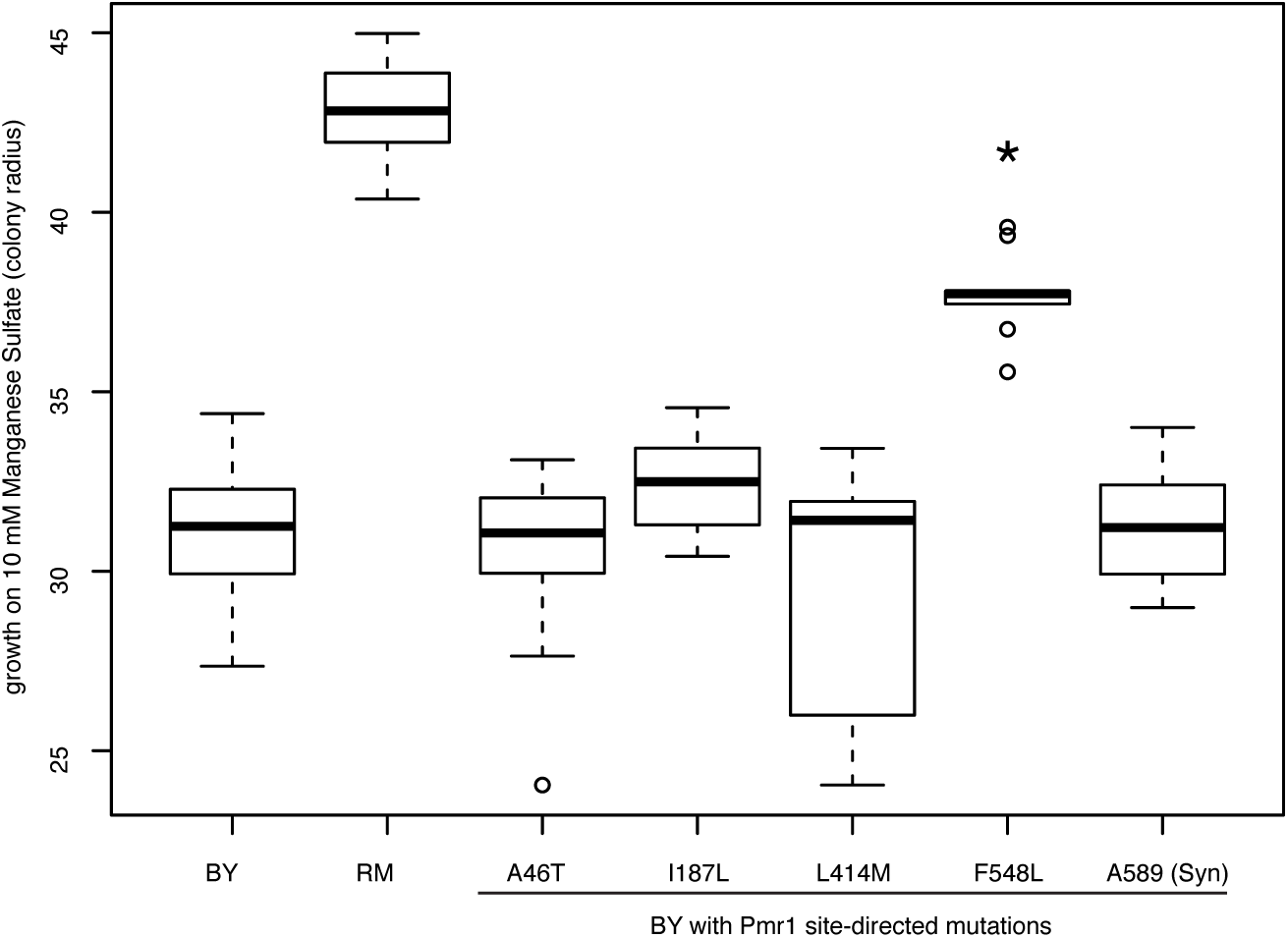
Direct introduction of Pmr1-F548L into BY enhances manganese resistance. oxplots of manganese sensitivity for strains with single *PMR1* variants introduced from RM into BY, along with the BY and RM parental strains (first and second leftmost boxes). n ≥ 10 for all genotypes. * p < 0.001 in comparison to BY, Welch's two-sided T-test.

We measured manganese sensitivity in this fine-mapping panel (Fig. 3B). Comparison of the panel phenotypes with the breakpoint locations pinpointed a single polymorphism as responsible for increased sensitivity in BY. The variant encodes a phenylalanine in BY and a leucine in RM at position 548 of Pmr1, a manganese transporter. Six lines had recombination events between Pmr1-F548L and the closest polymorphism to the right, 402 bp away, and were either fully sensitive or resistant to manganese, depending on which Pmr1-F548L allele was homozygous in the line. One line had a recombination between Pmr1-F548L and the closest polymorphism to the left, 125 bp away, and showed the intermediate manganese sensitivity phenotype expected for a heterozygote at the causal variant. LOD score analysis of the fine-mapping panel also identified a support interval containing only Pmr1-F548L (Fig. 3B).

To directly test the effect of Pmr1 variants on manganese sensitivity, we individually engineered into BY the RM alleles of Pmr1-F548L, the two neighboring polymorphisms, and the two remaining nonsynonymous Pmr1 polymorphisms, using a CRISPR-based variant replacement approach. As expected from the LOH fine-mapping, changing phenylalanine-548 to leucine conferred significant manganese resistance, whereas none of the other four polymorphisms had a significant effect (Fig. 4).

*PMR1* encodes an ion pump that transports manganese and calcium into the Golgi (Culotta *et al*. 2005). Pmr1 is a member of the P-type ATPase family of ion and lipid pumps found in all branches of life, and many other P-type ATPases have a conserved leucine at the position homologous to phenylalanine-548 of Pmr1. Solved structures of P-type ATPases with this leucine (Hilge *et al*. 2003) (Toyoshima and Mizutani 2004) show it directly contacting ATP (fig. S4). Furthermore, mutating the homologous leucine of the rabbit calcium pump to phenylalanine decreases its function by affecting ATP binding (Clausen *et al*. 2003). Thus, the F548L polymorphism is expected to reduce the ability of Pmr1^BY^ to transport manganese into the Golgi, relative to Pmr1^RM^, consistent with BY’s manganese sensitivity.

Pmr1 leucine-548 is conserved across fungi, with some species having an isoleucine or valine at the homologous position, and none with phenylalanine (fig. S5). In the *S. cerevisiae* population, almost all sequenced *PMR1* alleles have leucine-548, with phenylalanine-548 only in BY and other laboratory strains (Liti *et al*. 2009; Song *et al*. 2015) whose PMR1 alleles are likely directly related to BY (Schacherer *et al*. 2009). BY is derived from EM93, a diploid strain isolated from a fig (Mortimer and Johnston 1986). Sequencing of *PMR1* in EM93 revealed that EM93 is heterozygous for Pmr1-F548L (fig. S6), suggesting that either the mutation is not laboratory-derived or that it occurred between EM93’s isolation and its entry into a stock collection.

Decades of mapping studies have uncovered loci for myriad traits, but identification of the underlying genes and variants has lagged. Our CRISPR-assisted mapping approach promises to close this gap. In contrast to previous strategies, our method generates a higher density of recombination events, is easily targetable to any region of the genome, and does not require time-consuming extra generations of crossing to increase recombination frequency. Conversely, the strength of a traditional meiotic mapping panel is the ability to scan the entire genome. Complex traits, with multiple small-effect QTLs, pose a greater challenge for any mapping method. Importantly, in LOH mapping the rest of the genome outside the region targeted for LOH is held constant when a given QTL is being queried, thus effectively reducing the complexity of a trait by eliminating variance due to other segregating QTLs.

We anticipate that trait mapping with targeted LOH panels will aid efforts to understand the genetic basis of trait variation. In addition to applications in single-celled organisms, LOH panels could be generated from cultured cells, enabling in vitro genetic dissection of human traits with cellular phenotypes. In multicellular organisms, mapping resolution could be enhanced with CRISPR-directed meiotic recombination events. Indeed, the mutagenic chain reaction system developed in vivo in fruit flies (Gantz and Bier 2015) and mosquitos (Gantz *et al*. 2015; Hammond *et al*. 2015) uses CRISPR to generate gene conversion events in meiosis with high efficiency. Additionally, LOH in early development could generate chimeric individuals. The targeted LOH method also has the potential to be applied to viable interspecies hybrids that cannot produce offspring, allowing trait variation between species to be studied genetically beyond the few systems where it is currently possible (Orr and Presgraves 2000; Woodruff *et al*. 2010).

In addition to their research applications, targetable endonucleases hold promise for gene therapy (Hsu *et al*. 2014; Tebas *et al*. 2014). Certain disease alleles may be difficult to directly target by CRISPR because of their sequence complexity, such as the expanded trinucleotide repeats that underlie Huntington’s disease. In these cases, directing a DSB to occur in the vicinity of a pathogenic allele so that it is replaced with its nonpathogenic counterpart by LOH may represent a more feasible alternative.

## Acknowledgements

We thank Kruglyak laboratory members for helpful discussion, Steven Clarke for strain BY4742, George Church for plasmids, and Sri Kosuri for his flow cytometer. Funding was provided by the Howard Hughes Medical Institute and NIH grant R01 GM102308 (L.K.). Sequencing data was deposited at the Sequence Read Archive under accession SRP072527, and other data and code was deposited at https://github.com/joshsbloom/crispr_loh.

